# Non-redundant role of the α3 isoform of Na^+^,K^+^-ATPase in neuronal excitability and spiking dynamics

**DOI:** 10.1101/2025.10.24.684230

**Authors:** Kirill Reshetnikov, Vasilii Tiselko, Maxim Dobretsov

## Abstract

The Na^+^,K^+^-ATPase (NKA) plays a fundamental role in neuronal excitability by maintaining ionic gradients and contributing to electrogenic resting currents. Among its isoforms, the neuron-specific *α3* subunit exhibits a uniquely low affinity for intracellular Na^+^, weak voltage dependence, and slightly reduced ATP sensitivity compared to the ubiquitous *α1*. Although mutations in *ATP1A3* are known to cause severe neurological disorders, the specific kinetic features of *α3* that underlie its functional specialization remain incompletely understood. Here we employed biophysically detailed models of stretch receptor neurons, grounded in patch-clamp recordings of the *α3*-isoform current and spiking responses, to dissect the contribution of isoform-specific pump kinetics to firing behavior. Substitution of *α1* for *α3* abolished the ability to sustain long spike trains and reduced high-frequency entrainment, whereas *α3*-preserved prolonged discharges and faithful responses to vibratory stimuli. Remarkably, even halved pump density *α3*^50%^ preserved superior excitability compared to mixed expression (*α3*^50%^*/α1*^50%^), indicating that kinetic profile rather than pump quantity determines firing capacity. Hybrid models revealed that Na^+^ affinity is the decisive factor: retaining the low Na^+^ affinity of *α3* preserved excitability, while introducing *α1*-like voltage or ATP dependence produced only minor effect on simulated neuron discharge. These findings establish a mechanistic explanation for the selective expression of *α3* in muscle spindle afferents and other neurons with high-frequency demands, and help to explain why *α1* cannot compensate in *ATP1A3*-linked diseases. More broadly, they highlight the principle that isoform specialization of the NKA is not redundant but tuned to the discharge requirements of distinct neuronal populations.

## Introduction

Na^+^,K^+^-ATPase (NKA) is a complex of integral membrane protein subunits that sets and maintains Na^+^ and K^+^ gradients across the cell plasma membrane. As an electrogenic transporter, NKA also contributes to resting membrane potential and excitability set points, linking pump kinetics to firing behavior ^1^. The major α subunit of the mammalian enzyme has four isoforms (α isoforms of NKA) that are expressed in a cell type- and developmental stage-dependent manner. Thus, the neuronal *α3* NKA is abundant in nervous system but rarely detected in other tissues of adult vertebrates. In the CNS, the α1 NKA isoform is ubiquitously express across most cell types, whereas the neuronal α3 NKA is largely neuron-specific—it is abundant in the nervous system but rarely detected in other adult vertebrate tissues, and notably absent from glial cells ^2–5^. Furthermore, mapping studies show enriched *α3 NKA* expression within CNS is characteristic of specific neuonal populations, including in particular many GABAergic neurons in motor circuits ^2,3^. The reasons behind the existence and differential expression of multiple NKA isoforms – and specifically *α3* – remain unknown. However, genetic haploinsufficiency of *α3* results in severe neurological disorders with no signs of compensation by overexpression of other NKA isoforms ^6^. Clinically, ATP1A3 mutations underlie Rapid-Onset Dystonia Parkinsonism (RDP), Alternative Hemiplegia of Childhood (AHC) and related syndromes, and α1 upregulation does not happen as compensatory mechanism in neurons with insufficient *α3 NKA* ^6^. It is possible that non-uniform expression of NKA isoforms in the nervous system is associated with the existence of unique functional characteristics of the *α3* isozyme on one hand, and unique requirements for NKA activity imposed by the specific function and discharge characteristics of the neuron on the other. Multiple studies indicate the presence of several important kinetic features for *α3 NKA. Compared to other α isoforms of the transporter* ^1,7,8^, suggesting isoform-specific NKA operating regimes. Specifically, the majority of experiments in transfected cells suggest that compared to α1 and α2 isoforms, the *α3* NKA has about 3-fold lower affinity for activation by intracellular Na^+^, lower requirements for cellular ATP, and very limited – if any – activation of *α3 NKA* during cell depolarization (low voltage-dependence of the pump cycle). Reported values cluster around ∼10 mM for *α1* versus ∼30 mM for *α3*, consistent with selective recruitment of *α3* during activity-dependent Na+ loading ^1,7,9^. Combined, existing kinetic differences between isoforms of NKA make *α3* NKA specifically suitable for neurons whose normal function requires generation of prolonged, high-frequency discharges ^1,10^. In support of this idea, peripheral sensory systems, antibodies for *α3 NKA* selectively label plasma membrane of muscle spindle neurons ^10^, which are the only afferent neurons capable of sustaining prolonged, high-rate discharges and to follow vibration at high frequencies, whereas other primary afferents predominantly use the α1 isozyme(for several seconds at 25–100 action potentials/sec) ^11,12^. Nonetheless, it remains unknown, which of specific kinetic properties of *α3* NKA, listed above, play a decisive role in the firing capacity of spindle afferent neurons. To address this question here with *α3*- and *α1*-like pump kinetics (α3- and α1-neurons). In additional, to asses potential functional significance of haploinsufficiency of *α3* NKA and to isolate contribution of isoform differences in voltage and ATP dependence we studied models with 50% decreased *α3* NKA site density and constructed *α3* hybrids with *α1*-like voltage or ATP dependence with preserved all other ionic conductances. Our simulations were based on previously published computer simulation of a slowly adapting lobster stretch receptor neuron ^13^ modified to match the major passive properties of large dorsal root ganglion rat neurons ^14–16^ and supplied by NKA with characteristics known from previous studies for either *α3* or *α1* NKA ^1,7,8,17^. Discharge properties (activation threshold, response to maintained depolarization and rate of discharge frequency accommodation, and neural response to series of frequent stimuli) were compared between two simulated models and the results of physiological stretch receptor discharge studies ^18,19^.

## Results

### Properties of Na^+^,K^+^-ATPase isoforms: electrophysiology and model calibration

We investigated how Na^+^,K^+^-ATPase isoforms shape spiking dynamics using a biophysically detailed neuronal model. Isoform-specific pump kinetics were specified a priori, and the resulting firing behavior was examined. To ensure physiological plausibility, the model was anchored to electrophysiological recordings of the pump current obtained in dorsal root ganglion (DRG) neurons.

Figure 1a shows the steady-state holding current (*I*_*h*_) recorded from an isolated DRG neuron under conditions that stimulate the Na^+^/K^+^ pump. The external solution contained 140 mM Na^+^ and 5.4 mM K^+^; the patch-pipette solution contained 60 mM Na^+^ and 0 mM K^+^ (N-methyl-D-glucamine substitution). The holding potential (*V*_*h*_) was −40 mV (see Methods). After break-in (time zero in Fig. 1a), *I*_*h*_ increased in the positive (outward) direction due to activation *I*_*p*_ by Na^+^ entering the cell from the pipette. When *I*_*h*_ reached a steady state, 1 mM ouabain was applied (black bar above the trace in Fig. 1a) to block *I*_*p*_, producing a new steady state of *I*_*h*_. During washout, restoration of Na^+^/K^+^-pump activity was reflected by the return of *I*_*h*_ toward its pre-drug level. As in all similar patch-clamp experiments ^20–24^, *I*_*p*_ in our experiments was defined as the 1 mM ouabain–sensitive fraction of in *I*_*h*_ (arrow in Fig. 1a). Red symbols denote the average *I*_*p*_ obtained from 27 neurons under these conditions. As reported in other cell types^20,21^, *I*_*p*_ in DRG neurons exhibit time-dependent run-down; its rate was 2.0 ± 0.2% per min. To minimize the impact of run-down in subsequent analyses, *I*_*p*_ values were compared 10 min after establishment of the whole-cell configuration in each cell. These experiments demonstrate that DRG neurons exhibit a measurable *I*_*p*_ suitable for estimating Na^+^/K^+^-pump activity.

**Figure 1.**
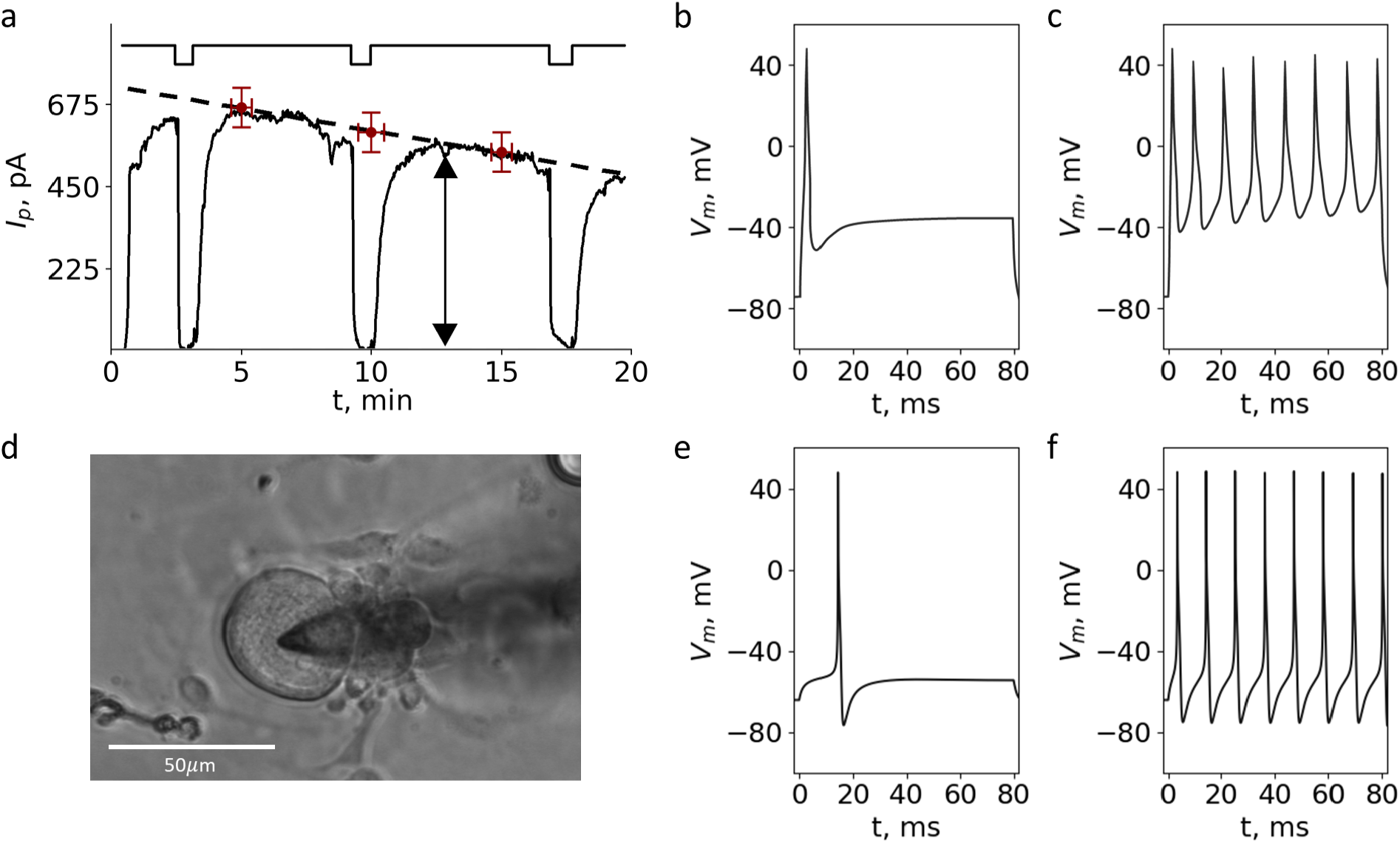
Electrophysiological calibration of Na^+^,K^+^-ATPase pump current and isoform-dependent effects on DRG firing. a. Steady-state holding current from an isolated DRG neuron under conditions that stimulate the Na^+^/K^+^ pump. After break-in (time 0), *I*_*p*_ increased in the outward direction as intracellular Na^+^ from the pipette activated outward pump current (*I*_*p*_). Application of 1 mM ouabain (bar) blocked pump current, shifting *I*_*p*_ to a new steady state; washout of ouabain resulted in restoration of the current toward the pre-drug level. Red symbols denote average IpI_pIp from 27 neurons. b,c. Representative current-clamp recordings illustrating contrasting responses to a depolarizing step: a neuron producing a single/short-burst response (b) versus a neuron that fires continuously throughout the step (c); the continuously firing neuron expresses the α3 Na^+^,K^+^-ATPase isoform. D. Patch pipette is approaching isolated DRG neuron from the right. e,f. Model simulations incorporating isoform-specific pump kinetics reproduce the experimental differences in firing: simulated pump currents and the resulting spiking behavior recapitulate the characteristic divergences observed in recordings (f,g).

Studies in intact rat DRG have identified a subpopulation of large neurons that fire continuously throughout a depolarizing voltage step; relative to neurons that produce a single spike or a short burst, this persistent firing was suggested to be linked to specific properties of the Na^+^,K^+^-ATPase in these cells^25^. In our patch-clamp experiments, we likewise encountered DRG neurons with such behavior (Fig. 1b,c,d), in which a prolonged spiking response in an α3-isoform–expressing neuron displayed qualitatively different spiking dynamics. Figures 1d shows example of a patch-clamped DRG neuron. By incorporating the properties of distinct Na^+^,K^+^-ATPase isoforms, our model reproduced differences in spiking responses consistent with these experimental observations (Fig. 1e,f); specifically, the simulated pump currents recapitulated the characteristic divergences in firing seen in the electrophysiological records.

In all simulations (Fig. 2), pump current for α1 and α3 was parameterized with experimentally grounded dependencies on membrane voltage, intracellular Na^+^, and ATP (see ^1^ and Methods). Specifically, Na^+^ affinity was set to reflect the established isoform difference—half-maximal activation (K_0_._5_) of ∼10 mM for *α1* versus ∼30mM for *α3* (Fig. 2A). Voltage dependence was implemented so that *α3* exhibited weaker voltage sensitivity than *α1* (Fig. 2B). Finally ATP dependence was modeled such that at physiological ATP (∼3 mM) *α1* operated near saturation (∼98% *I*_*p*,,*max*_) and *α3* slightly below (∼95% *I*_*p*,,*max*_) (Fig. 2C). These specifications capture the prevailing view that *α1* NKA is strongly active at low [Na^+^]_i_, whereas *α3* is preferentially recruited as [Na^+^]i rises during sustained activity^1^.

**Figure 2.**
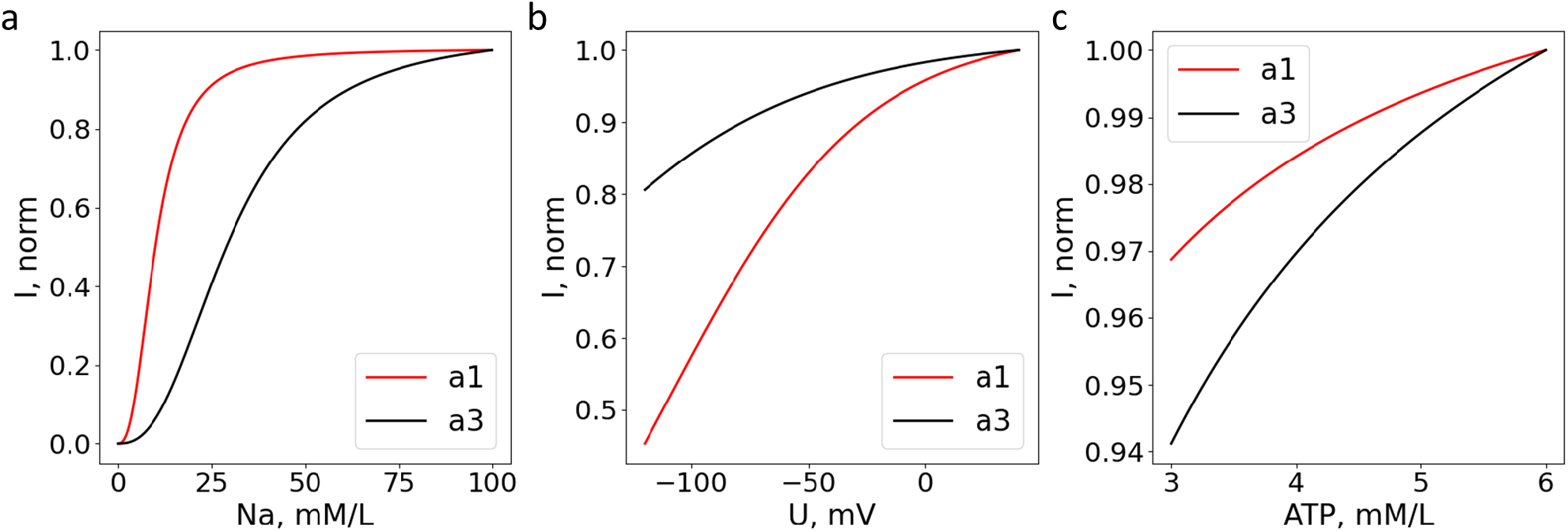
Isoform-specific pump kinetics implemented in the model. (a) Steady-state pump activity as a function of intracellular Na^+^ concentration ([Na^+^]i) for α1 (red) and α3 (black). Curves are normalized to each isoform’s maximal turnover rate. Half-maximal activation (K_0_._5_) is ∼10 mM for α1 and ∼30 mM for α3, mimicking lower apparent Na^+^ affinity of α3 relative to α1. (b) Voltage dependence. Normalized pump activity versus membrane potential (U). Hyperpolarization suppresses α1 more strongly than α3; at U ≈ –100 mV, α1 operates at ∼50% of *I*_*p*,*max*_ whereas α3 retains ∼80% of *I*_*p*,*max*_. (c) ATP dependence. Normalized pump activity versus intracellular ATP. Both isoforms are near-saturated at physiological ATP (∼3 mM), with α1 slightly closer to saturation (∼98% *I*_*p*,*max*_) than α3 (∼95% *I*_*p*,*max*_). These empirically constrained dependencies were used a priori to parameterize isoform-specific Na^+^/K^+^-ATPase kinetics in all simulations (Methods). The resulting equilibrium set points for each condition are reported in Table 1. In this study, we focus on modeling the spiking dynamics of large DRG stretch neurons of rats. To this end, we consider large-diameter cells characterized by a predominant expression of the α3 isoform. The maximal Na^+^/K^+^ pump current was accordingly calibrated for large cells based^17^, and the resulting steady-state parameters are summarized in Table 1. XXX

Notably, despite the stronger voltage-dependent suppression of *α1* at very negative potentials, at resting U and low-to-moderate [Na^+^]i its higher Na^+^ affinity yields a larger net pump current; accordingly, the steady-state values reported in Table 1 are the model’s equilibrium set points under these kinetic assignments. Throughout, *α3*^50%^ denotes half-density expression of the *α3* pump (i.e. 50% of baseline *α3*) without compensatory substitution by other isoforms and with identical *α3* kinetics, whereas *α3*^50%^/*α1*^50%^ denotes co-expression of 50% *α1* and 50% *α3*, each retaining its native kinetic profile.

**Table 1.** Resulting baseline state variables for each pump configuration used in the simulations. Resting membrane potential (V_rest, mV), intracellular ion concentrations ([K^+^]i and [Na^+^]i, mM), and steady outward Na^+^/K^+^-ATPase current (“Pump”, pA) obtained after model equilibration under no-stimulus conditions. *α3*^50%^ denotes half-density expression of *α3* with identical *α3* kinetics and no compensatory substitution by other isoforms; *α3*^50%^/*α1*^50%^ denotes co-expression of 50% *α1* and 50% *α3*, each retaining its native kinetic profile. *α3*^VD^ and *α3*^ATP^ are *α3*-based hybrids in which only the indicated voltage and ATP dependencies were replaced by *α1*-like characteristics while retaining *α3* Na^+^ affinity. All other membrane and channel parameters were identical across conditions. Positive pump current denotes outward (hyperpolarizing) electrogenic flux. Values represent the model’s resulting steady-state set points under the specified kinetic assignments.

| Na<sup>+</sup>/K<sup>+</sup>-ATPase isoforms | V<sub>rest</sub>, mV | [Na<sup>+</sup>]<sub>i</sub>, mM | [Na<sup>+</sup>]<sub>o</sub>, mM | I<sub>pump,max</sub>, pA |
| --- | --- | --- | --- | --- |
| $\alpha 1$ | -65 | 147 | 7 | 800 |
| $\alpha 3$ | -62.5 | 130.5 | 22 | 786 |
| $\alpha 3^{50\%}$ | -60. | 117 | 35 | 752 |
| $\alpha 3^{50\%}/\alpha 1^{50\%}$ | -65 | 146.25 | 10.25 | 790 |
| $\alpha 3^{VD}$ | -65 | 132 | 20 | 782 |
| $\alpha 3^{ATP}$ | -65 | 131.5 | 20.5 | 779 |

**Spike Frequency Adaptation and Input–Output Response with Different Pumps**

We first examined how pump kinetics impact neuronal excitability. Under a prolonged depolarizing step current (3.5 s), the *α3*-model neuron sustained spiking for the entire stimulus, with only a gradual slowing of spike frequency. In stark contrast, the *α1* neuron produced an initial high-frequency burst but ceased firing after ∼1–2 s despite continued stimulation; the membrane then settled at a subthreshold potential. Notably, the 50%-*α3* condition (half the normal *α3* pump density) fired continuously throughout the stimulus and at even higher initial frequency, exhibiting weaker spike-frequency adaptation than the full *α3* model (Fig. 3B, Fig. 4A).

**Figure 3.**
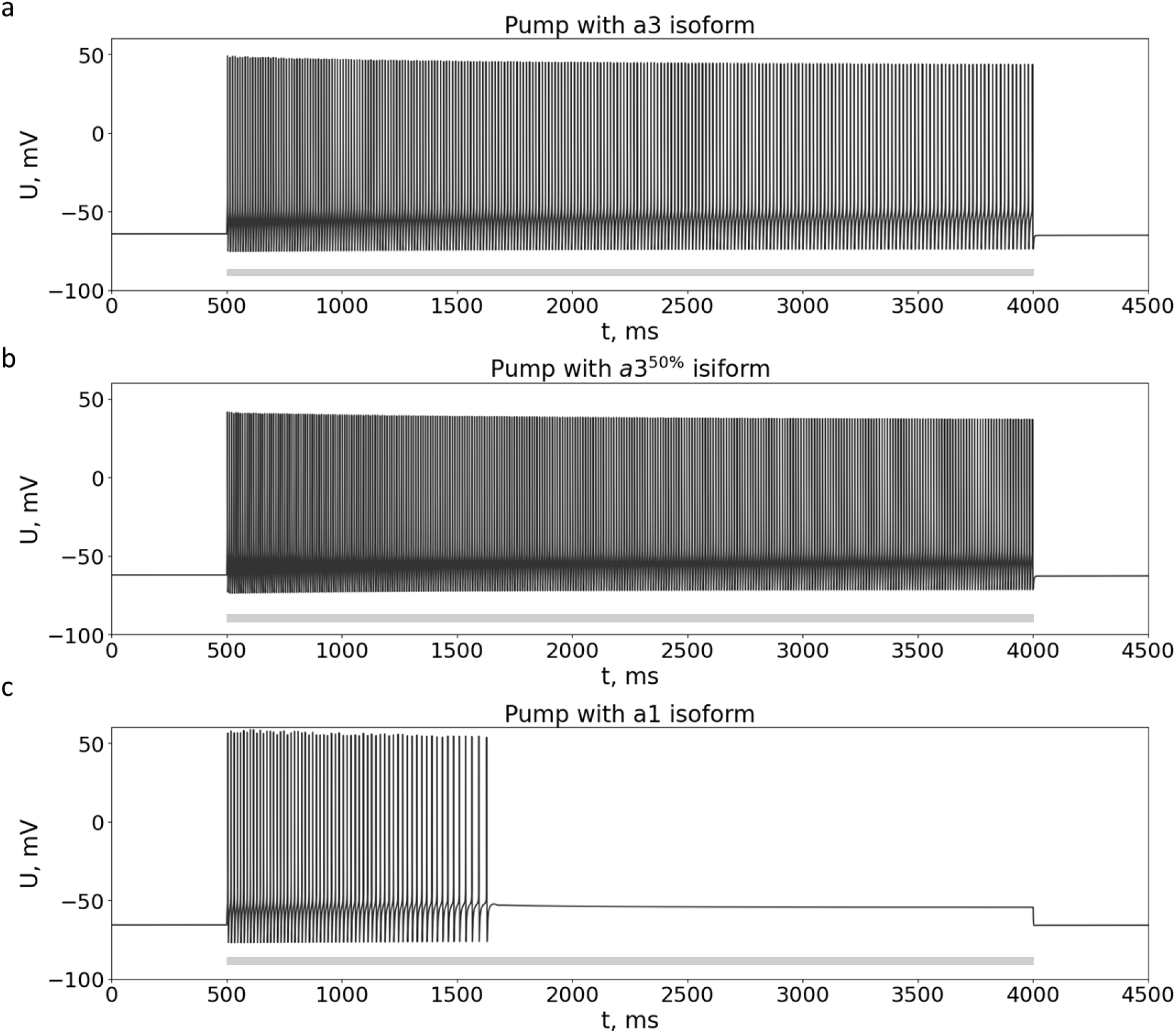
Spike-train responses to a prolonged depolarizing step under different pump isoform conditions. Spike trains are shown for (a) α3, (b) 50% α3 (half-density α3 with identical kinetics and no substitution by other isoforms), and (c) α1. The grey bar beneath each trace indicates the current-step stimulus epoch (identical amplitude and duration across panels; 3.5 s rectangular step beginning at t = 1 s). All other membrane and channel parameters are identical across conditions; simulations start from the condition-specific steady state (Table 1). (a) α3: continuous spiking persists for the full 3.5 s step with moderate spike-frequency adaptation (gradual rate decline over time). (b) 50% α3: spiking also persists throughout the step but with a higher initial frequency and the weakest adaptation among the conditions (shallower decay of instantaneous rate than in α3).(c) α1: pronounced adaptation leads to early failure; the spike train terminates ∼1 s after stimulus onset despite ongoing current injection, and the membrane potential stabilizes at a subthreshold level. Together, these panels illustrate that *α3* kinetics—even at half density—support sustained high-rate firing under tonic drive, whereas *α1* kinetics precipitate rapid spike-frequency adaptation and loss of spiking.

**Figure 4.**
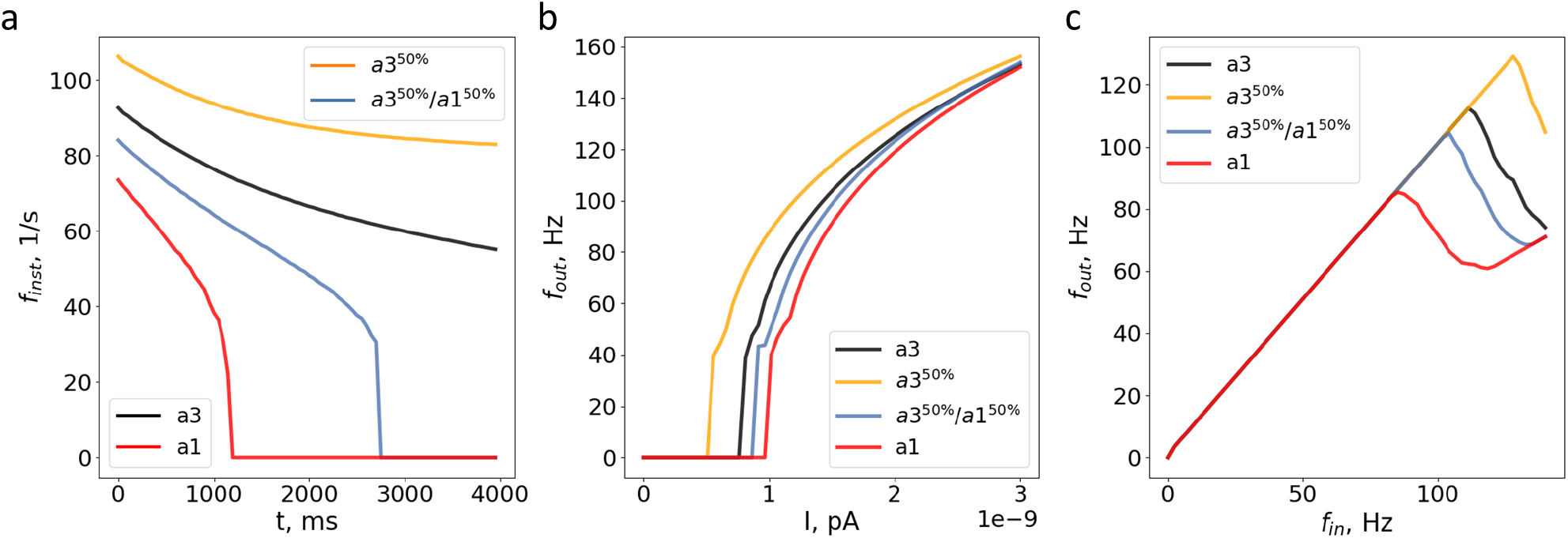
Comparative firing dynamics across pump isoform conditions. All models used identical membrane/channel parameters and were initialized from the condition-specific steady state (Table 1). (a) Spike-frequency adaptation during a sustained step. Instantaneous firing frequency (from interspike intervals) over a 4-s depolarizing step. *α1* shows rapid decay and early failure (∼1 s); mixed model *α3*^50%^/*α1*^50%^ also fails to fire after about 2 s of activity. Thereas *α3* model maintains a continuous firing with moderate adaptation for entire duration of stimulus. The *α3*^50%^ model sustains the highest rates with the weakest adaptation. (b) Frequency–current (f–I) relations. Steady-state firing rate versus injected current, measured over the last 1 s of a 4-s step for each amplitude. Rheobase is lowest for *α3*^50%^, slightly higher for *α3*, and higher for mixed *α3*^50%^/*α1*^50%^ and *α1*, which are similar to each other. Above threshold, all curves rise and then shallow, with isoform-dependent threshold shifts accounting for most differences. (c) Entrainment to periodic stimulation. Output firing frequency versus input pulse frequency for trains of 1-ms current pulses (fixed amplitude). All conditions follow 1:1 at low frequencies; the loss of 1:1 occurs earliest for *α1*, then for *α3*^50%^/*α1*^50%^, later for *α3*, and latest for *α3*^50%^. Across paradigms, performance ranks as: most pronounced firing response *α3*^50%^, next *α3*, then *α3*^50%^/*α1*^50%^, and lowest *α1*.

Thus, replacing the high-affinity, strongly voltage-dependent *α1* pump with *α3* dramatically improves the ability to fire repetitively under sustained neuron depolarization (mimicking sustained muscle stretch). Even at a reduced density of *α3* NKA neuron outperforms purely *α1 NKA neuron*.

Mechanistically, *α3*’s low Na^+^ affinity delays strong pump engagement until Na^+^ has risen during spiking; once engaged, *α3* clears Na^+^ loads effectively without prematurely curtailing excitability. In contrast, *α1*’s high baseline activity (due to its higher Na^+^ affinity and voltage sensitivity) causes earlier and stronger pump activation, accelerating spike-frequency adaptation and leading to firing failure.

To quantify firing dynamics, we measured instantaneous spike frequency over time during a sustained current step and constructed spike-frequency adaptation (SFA) curves (Fig. 4A). The *α1* neuron showed steep SFA and early spiking failure: firing frequency peaked around 70 Hz and fell to 0 Hz within ∼1 s. The mixed condition *α3*^50%^/*α1*^50%^ (co-expression of 50% *α1* and 50% *α3*) also failed, dropping to 0 Hz by ∼3 s. In contrast, the *α3* neuron maintained a continuous spike train, with frequency decaying from ∼90 Hz to ∼60 Hz by 4 s. The *α3*^50%^ neuron exhibited the least adaptation, declining only from ∼110 Hz to above 80 Hz by 4 s.

Hence, exclusive *α3* expression—even at half density—outperforms *α3*^50%^/*α1*^50%^, underscoring that kinetic profile outweighs pump quantity for sustaining activity. The inability of *α3*^50%^/*α1*^50%^ to maintain firing aligns with in vivo observations that neurons relying on *α3* do not compensate by upregulating *α1* ^6^. In other words, even partial *α1* engagement imposes an electrogenic load that curtails excitability, masking the benefits of *α3*.

The *α3* pump endowed the neuron with a lower rheobase (threshold current) and a higher sustainable firing rate compared to *α1* (Fig. 4B). The *α1* model required a stronger input current to initiate spiking (rheobase ∼20% higher than in *α3*) and its firing rate saturates at a modestly lower level. The *α3*^50%^/*α1*^50%^ closely tracked *α1* (elevated threshold and reduced dynamic range). Meanwhile, the *α3*^50%^ neuron retained a low threshold comparable to full *α3* and achieved high firing rates (∼120 Hz at strong input), nearly matching the full *α3* model. Thus, *α3* kinetics expand the input–output dynamic range by maintaining ionic homeostasis at high activity while avoiding excessive pump drive at rest.

Resting set-points differed systematically across isoform conditions and parallel the f–I results (see Table 1). *α1*-expression neurons were more hyperpolarized (∼–65 mV resting potential) with low resting [Na^+^]i (∼7 mM) and a larger resting pump current (≈800 pA outward). In contrast, *α3* and *α3*^50%^ neurons rested at less hyperpolarized potentials (∼–63.5 and –61.4 mV, respectively) with higher [Na^+^]i (18.5 and 27.5 mM) and smaller pump currents (∼798 and 779 pA). The *α3*^50%^/*α1*^50%^ (–65 mV; ∼790 pA) mirrored the *α1*-like state. A lower baseline electrogenic load and higher internal Na^+^ under *α3* shift the neuron toward easier excitability (lower rheobase), whereas *α1*’s stronger resting pump activity makes the neuron less excitable (higher rheobase). The *α3*^50%^/*α1*^50%^ mixed condition’s *α1*-like resting parameters explain its *α1*-like firing threshold.

Performance in sustained firing scaled inversely with the resting pump current: the best performer *α3*^50%^ also had the smallest pump current at rest (∼779 pA), whereas *α1* and *α3*^50%^/*α1*^50%^, with the largest pump currents (∼800 and 795 pA), showed the earliest failure. Thus, *α3*’s kinetic profile provides high-frequency firing capacity at a lower baseline energetic burden.

Across all measures—adaptation over time (Fig. 4A), steady-state f–I dynamic range (Fig. 4B), and high-frequency entrainment (Fig. 4C, described next)—the rank order of performance was consistent: best *α3*^50%^, next *α3*, then *α3*^50%^/*α1*^50%^, and lowest *α1*. This cross-paradigm consistency strengthens the conclusion that isoform kinetics, rather than pump abundance per se, determine the neuron’s spiking capacity. Given the weak voltage suppression of *α3* (Fig. 2B) and its low Na^+^ affinity (Fig. 2A), pump activity is minimal at rest and early in a spike train, but rises as [Na^+^]i accumulates. During repetitive firing, *α3* remains sufficiently active even through afterhyperpolarizations (AHPs) to clear Na^+^ without over-hyperpolarizing the membrane, thereby stabilizing interspike trajectories and preventing depolarization block. By contrast, the *α3*^50%^/*α1*^50%^ model behaved functionally like the *α1*-only model in both paradigms. This non-additivity indicates dominance of the high-affinity *α1* pathway near threshold: even a partial *α1* component imposes an outward pump current sufficient to curtail excitability, negating the benefits of *α3* co-expression.

### High-frequency spiking responses to vibratory stimuli

A key feature for stretch receptor neurons is the ability to follow rapid periodic inputs with ∼1 spike per cycle. When driven with brief 1-ms pulses at increasing frequencies (Fig. 4C), the *α3* model preserved 1:1 activation up to ∼110–120 Hz. In contrast, the *α1* neuron lost 1:1 activation much earlier (around 80 Hz), producing skipped spikes as the input frequency increased.

The *α3*^50%^/*α1*^50%^ neuron again resembled the *α1*-only case, losing faithful 1:1 following just above 100 Hz. The *α3*^50%^ neuron matched or slightly exceeded the full *α3*, maintaining 1:1 spiking up to ∼120+ Hz. This superior high-frequency following is consistent with its weaker adaptation and lower baseline electrogenic load, while still preserving *α3*’s robust Na^+^ clearance under high loads. Overall, exclusive *α3* expression is necessary for reliable high-frequency encoding, whereas any substantial *α1* component undermines fast entrainment.

### Contributions of specific pump kinetic features to firing behavior

Finally, we isolated the contributions of voltage dependence and ATP dependence (aside from Na^+^ affinity) by implementing hybrid pump models. These hybrids were based on the *α3* isoform but with either the voltage dependence or the ATP sensitivity replaced by the *α1*-like characteristic (Fig. 5), while retaining *α3*’s low Na^+^ affinity in both cases.

**Figure 5.**
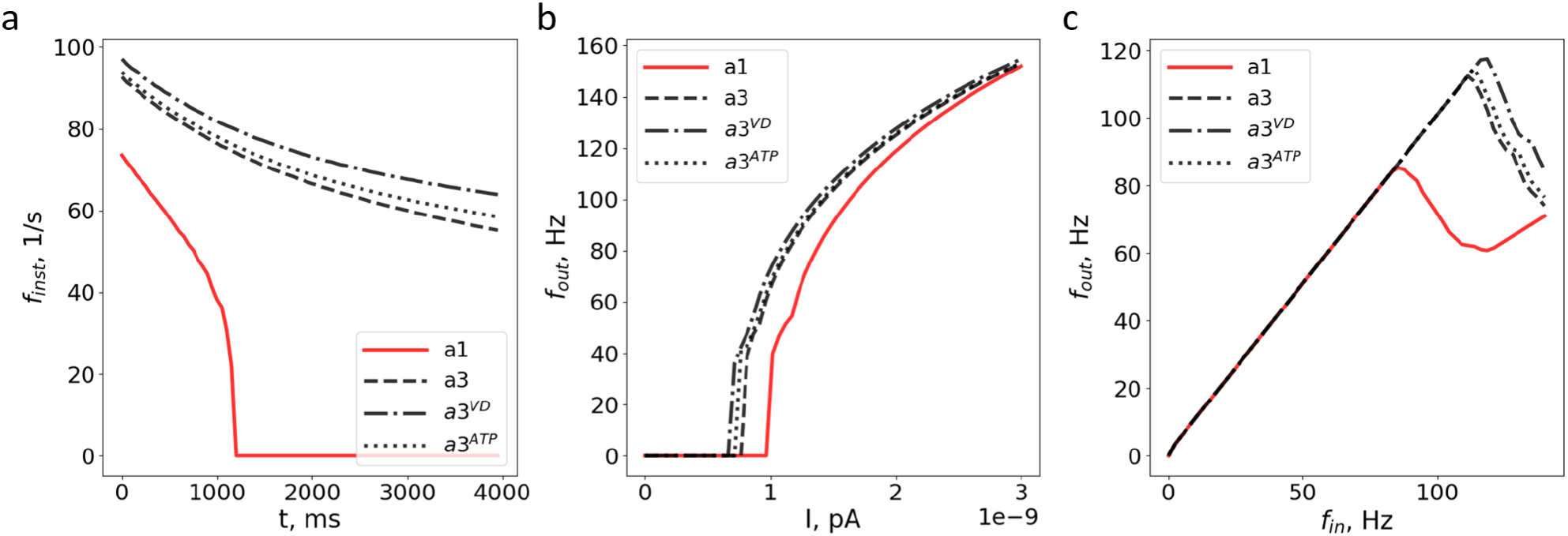
Mechanistic substitutions within *α3* isolate the roles of voltage and ATP dependence. Legend: *α1* (red), *α3* (black), *α3*^VD^ *(α3* with *α1*-like voltage dependence, dashed), *α3*^ATP^ (*α3* with *α1*-like ATP dependence, dot–dash). Both hybrids retain *α3*-like Na^+^ affinity; only the indicated dependency was replaced. Membrane/channel parameters are identical across conditions. Simulations start from the condition-specific steady state (Table 1). (a) Spike-frequency adaptation during a sustained step. Instantaneous firing frequency over a 4-s depolarizing step of fixed amplitude. *α1* shows rapid decay and early failure (∼1 s). *α3* maintains a continuous train with moderate adaptation. The *α3*^VD^ hybrid exhibits the weakest adaptation (trajectory highest among *α3*-like curves), while the *α3*^ATP^ hybrid closely tracks *α3* but remains slightly higher throughout. (b) Frequency–current (f–I) relations. Steady-state firing rate versus injected current, measured over the last 1 s of a 4-s step. The *α3*^ATP^ hybrid is virtually indistinguishable from *α3* in rheobase and gain. The *α3*^VD^ hybrid shows a slightly lower rheobase (left-shift) and reaches saturation at a somewhat lower current level. *α1* requires the highest threshold and saturates earlier at lower rates. (c) Entrainment to periodic stimulation. Output firing frequency versus input pulse frequency for trains of 1-ms current pulses (fixed amplitude). Loss of 1:1 entrainment occurs earliest for *α1*, later for *α3*, later still for the *α3*^ATP^ hybrid, and latest for the *α3*^VD^ hybrid. Across panels, substituting *α1*-like voltage or ATP dependence into *α3* produces only modest shifts relative to *α3*, whereas the full *α1* phenotype shows early failure and reduced fidelity. Preserving *α3*-like Na^+^ affinity maintains sustained firing and high-frequency following; voltage and ATP kinetics act primarily as secondary modulators of threshold, adaptation, and entrainment.

Both hybrid pumps exhibited spike-frequency adaptation almost as in the native *α3* model, with only slight reductions in adaptation compared to *α3*. The *α3*^VD^ hybrid showed the weakest spike-frequency adaptation of all *α3*-based models; its firing frequency trajectory remained a few Hz above the native *α3* throughout the stimulus. Mechanistically, the stronger *α1*-like voltage inhibition (greater pump activity suppression at hyperpolarized voltages) means the pump is less active during AHP/subthreshold periods, slightly lowering the baseline outward current without compromising high-load Na^+^ clearance (since the Na^+^ affinity is still *α3*-like). The *α3*^ATP^ hybrid nearly overlapped the native *α3* in firing pattern, but its frequency was consistently marginally higher (indicating marginally weaker adaptation). Given that intracellular ATP is near-saturating under physiological conditions, the effect of swapping ATP sensitivity was very small. The *α3*^ATP^ hybrid was virtually indistinguishable from native *α3* in rheobase and gain, confirming that the slight difference in ATP dependence does not affect excitability within normal ATP levels. The *α3*^VD^ hybrid exhibited a slightly lower rheobase (a left-shift of the f–I curve) relative to native *α3*, consistent with its reduced pump activity at subthreshold voltages and thus reduced initial outward current load.

Both hybrids extended the 1:1 following limit beyond that of the native *α3*. The *α3*^ATP^ hybrid maintained 1:1 entrainment to slightly higher frequencies than *α3*, whereas the *α3*^VD^ hybrid had the highest 1:1 cutoff frequency of the three, in line with its particularly weak adaptation profile.

Notably, neither single-property substitution reproduced the severe firing failure seen with the full *α1* pump. This indicates that the decisive determinant of firing behavior is the *α3*-like low Na^+^ affinity; differences in voltage or ATP dependence act as secondary modulators that fine-tune the threshold, adaptation, and entrainment without altering the qualitative ability to sustain firing. If *α3*’s Na^+^ affinity were hypothetically increased to *α1*-like levels, the pump would engage too early and too strongly at modest [Na^+^]i, likely recreating the *α1*-type rapid adaptation and spike failure. In summary, the *α3* isoform’s kinetic profile—especially its low Na^+^ affinity combined with limited voltage-induced suppression—underlies the capacity of stretch receptor neurons to sustain high-rate spiking and to faithfully encode rapid periodic inputs. Mixed or partial expression of *α1* cannot replicate this profile, which reinforces why these neurons predominantly rely on *α3* in vivo ^6^.

## Discussion

### Sustained Firing and Vibration Encoding in Mechanoreceptors

One of the most valuable outcomes of *α3* expression in our model neuron was the ability to maintain prolonged, high-frequency spike trains under sustained or vibratory stimuli. With *α3*, the model neuron fired continuously for the entire 3.5 s sustained stimulus at high frequencies, whereas the *α1*-expressing neuron rapidly adapted and fell silent after ∼1–2 s. Importantly, even the *α3*^50%^ model outperformed a full complement of *α1* pumps, supporting the idea that qualitative kinetic traits trump quantitative pump capacity. These findings directly address the longstanding question of why certain neurons (like muscle spindle afferents) uniquely express the *α3* isoform ^1,10^. Muscle spindle primary endings are known for their ability to fire at high rates for prolonged periods, encoding muscle stretch and especially high-frequency vibrations. Classic microneurographic studies in humans and cats showed that tendon vibrations selectively drive primary (Ia) spindle afferents into 1:1 phase-locked firing at vibration frequencies up to ∼100 Hz or more ^18,26^. In cats, most primary endings can follow sinusoidal stretches up to 150–200 Hz (and some even 300–500 Hz at sufficient amplitude) with one spike per cycle ^26^. By contrast, secondary spindle afferents and Golgi tendon organs cannot sustain such high-frequency tracking ^18^. Our results provide a biophysical explanation: the *α3* pump endows neurons with a broader firing bandwidth by delaying Na^+^-pump engagement until Na^+^ load is high. In a neuron built for endurance and speed – such as a primary spindle afferent – *α3* allows repetitive spikes without the premature hyperpolarizing feedback that an *α1* pump would impose. In essence, *α3* keeps the “brakes” off during the initial phase of intense activity, only ramping up Na^+^ extrusion once it’s absolutely necessary. This prevents early spike-frequency adaptation and helps avert depolarization block, thereby preserving 1:1 stimulus–response coupling at high frequencies.

Anatomical and molecular surveys report that neuron classes with high firing demands preferentially express *α3* and often exclude *α1*, and in disease or genetic perturbations of *α3*, compensatory upregulation of *α1* is generally not observed in the affected neurons ^3,6^. The selective expression of *α3* in muscle spindle afferents is strongly supported by histological and comparative data. Dorsal root ganglion neurons that innervate muscle spindles show intense *α3* immunoreactivity, whereas other large-fiber sensory neurons (e.g. cutaneous touch or joint receptors) do not ^3,10^. In fact, *α3* immunostaining has proven to be a selective marker for functionally identified spindle afferents ^10^. This pattern is conserved across species: the *α3* isoform is uniquely expressed in both the afferent (Ia fiber) and efferent (γ-motoneuron) neurons of the muscle spindle circuit ^27^. Such phylogenetic preservation of *α3* in spindle-related neurons underscores its critical role. Our modeling results align well with this specialization. They suggest that without *α3*, a spindle afferent would likely fail to faithfully encode high-frequency muscle vibrations, undermining its role as a vibration sensor in proprioception. Similarly, γ-motoneurons – which must fire tonically to maintain spindle sensitivity – likely rely on *α3* to sustain steady firing without fatiguing. In these neurons, an *α1*-dominated pump would cause excessive early adaptation, reducing the ability to hold the spindle taut during prolonged postural or vibratory stimuli.

It is noteworthy that our model was originally developed for a slowly-adapting stretch receptor neuron, yet with *α3* it exhibited response properties more akin to a rapidly-adapting Ia afferent. Slowly adapting spindle afferents (type II) typically have low sensitivity to high-frequency vibration and do not show 1:1 driving beyond ∼50–100 Hz ^18^. In contrast, the primary Ia fibers (which are *α3*-positive) excel at high-frequency following ^26^. The ability of our *α3* model to maintain firing up to ∼120 Hz and beyond suggests that isoform kinetics can effectively convert a neuron from a phasic “burst-and-fail” responder into a tonic high-frequency encoder. Real primary endings achieve even higher rates than our simulation, likely because in vivo they benefit from specialized morphology (multiple encoding sites along the intrafusal fiber) and modulatory fusimotor input. For instance, the primary ending has been described as a sensory organ with pacemaker-like properties stemming from distributed transduction sites that can each trigger spikes ^28,29^. This architecture, combined with γ-motor drive, can support baseline rhythmic firing and prolong the dynamic range of responses ^30^. We speculate that *α3* pump kinetics complement these morphological features: by keeping the resting pump current low, *α3* may facilitate spontaneous or low-threshold firing (pacemaker activity) in the absence of stimuli, and by engaging gradually, it prevents the cumulative Na^+^ influx from silencing the receptor during prolonged high-frequency activation. In summary, the *α3* isoform appears to be a molecular adaptation for endurance and fidelity in sensory neurons that must continuously encode rapid or long-lasting stimuli.

### Clinical Significance: α3 Na/K-ATPase in Neurological Disorders

The distinctive kinetics of the *α3* pump are not only vital for sensory encoding but also for normal function of many CNS circuits – a fact highlighted by the severe neurological disorders caused by ATP1A3 mutations. Heterozygous loss-of-function or missense mutations in the *α3* Na^+^/K^+^-ATPase underlie diseases such as rapid-onset dystonia–parkinsonism (RDP), alternating hemiplegia of childhood (AHC), and related syndromes ^31^. These conditions are characterized by episodic or progressive motor dysfunction, including dystonic spasms, parkinsonian features, seizures, hemiplegic attacks, ataxia, and cognitive impairment ^32,33^. Our findings shed light on why neurons are so sensitive to *α3* dysfunction. We saw that *α3*^50%^/*α1*^50%^ expression (or partial *α3* function) in a model neuron did not preserve the *α3* firing phenotype; instead, the high-affinity *α1* activity dominated, causing failure of sustained firing. This lack of functional compensation by *α1* mirrors what is observed in vivo. Neurons that normally rely on *α3* do not simply upregulate *α1* when *α3* is deficient ^3,6^. For example, cerebellar Purkinje cells express *α3* at very high levels and essentially no *α1* ^6^. Consequently, in *α3* mutant mice or upon selective *α3* inhibition, Purkinje cells cannot maintain normal firing, leading to cerebellar dysfunction and dystonia ^34^. Similarly, certain brainstem and spinal neurons lack an *α1* backup and are severely affected by *α3* loss ^33^. It has been suggested that γ-motoneurons (spindle efferents) in the spinal cord predominantly express *α3* and not *α1*, making spinal sensorimotor circuits particularly vulnerable in ATP1A3 disorders ^33^. This could manifest as impaired muscle spindle regulation and hence abnormal muscle tone or reflexes, although overt stretch-reflex deficits have not been well documented clinically in conditions like RDP or AHC. Given our simulation results, one might predict subtle proprioceptive abnormalities – for instance, reduced vibration sensitivity or impaired reflex modulation – in patients with *α3* haploinsufficiency. These may be overshadowed by the more prominent central symptoms (dystonia, paralysis episodes), but they warrant investigation.

A unifying hypothesis for ATP1A3-related diseases is that loss of *α3* pump function upsets the delicate excitation–inhibition balance in the brain ^6,35^. The *α3* isoform is heavily expressed in many fast-spiking and high-workload neurons, notably GABAergic inhibitory interneurons in the cortex, hippocampus, basal ganglia, and cerebellum ^2,36^. Immunohistochemical mapping showed intense ATP1A3 labeling in GABAergic neurons of movement control centers – striatum, globus pallidus, subthalamic nucleus, cerebellar cortex – consistent with the motor symptoms of RDP ^2^. In hippocampus and cortex too, most *α3*-positive cell bodies are GABAergic ^36^. These inhibitory neurons typically fire high-frequency trains to restrain circuit excitation (for example, cerebellar Purkinje cells tonically inhibit deep nuclei, and pallidal neurons provide constant inhibitory output). If *α3* function is impaired, our results predict that such neurons would struggle to sustain firing or would adapt too quickly. The outcome would be a net reduction in inhibitory output and thus network disinhibition. Indeed, patients with ATP1A3 mutations often exhibit hyperkinetic movements (dystonia, episodic spasms) and lowered seizure thresholds, which are hallmarks of diminished inhibitory control in motor and cortical circuits ^32,33^. Additionally, *α3* is expressed in some excitatory neurons that have high firing demands (for instance, certain glutamatergic neurons in thalamus and cortex), so *α3* loss can also directly weaken those neurons’ ability to sustain activity necessary for normal function (potentially contributing to paralysis attacks in AHC or cognitive deficits) ^31^. In short, the kinetic advantages of *α3* that we identified – support of repetitive spiking without failure – are precisely what is lost in ATP1A3 disorders, leading to neurons that cannot keep up with physiological demand.

Another aspect our study illuminates is the energy-use strategy of different isoforms. At physiological ATP concentrations, *α1* operates near its maximal rate whereas *α3* is slightly further from saturation ^37^. While we found ATP affinity differences had only minor firing effects, the implication is that *α3* may be a bit less “eager” for ATP and thus leaves a small reserve capacity. Coupled with its low resting Na^+^ turnover, an *α3*-dominant neuron has a lower baseline energy expenditure (in our model, the *α3*^50%^ neuron had the smallest resting pump current and presumably lowest ATP consumption). Neurons with high *α3* expression might therefore be less prone to energy depletion during intense activity, at least in theory. In pathological states, however, if *α3* is partially impaired, neurons might become metabolically stressed as they attempt (via *α1* or remaining *α3*) to restore ion gradients. Some ATP1A3 mutations indeed perturb ion binding and pump cycling ^34,38^, which could reduce pump efficiency and exacerbate energy strain. The clinical observation that episodes in RDP/AHC are often triggered by stress, energy demand, or excitatory challenges (fever, exercise, emotional stress) ^34^ aligns with the idea that neurons on the edge of pump failure decompensate when pushed. Our model’s behavior under partial *α3* function (mixed isoforms) – performing acceptably at low firing rates but failing under sustained load – is a simplified parallel to how patients remain largely normal until a stressor precipitates an episode. Furthermore, the fact that *α1* does not compensate for *α3* loss ^39^ underscores an important conclusion: it is not the quantity of pumping that matters, but the kinetic tuning of the pump to the task. This is why gene therapy or pharmacological strategies for ATP1A3 diseases are exploring ways to restore *α3* function specifically, rather than boosting general pump activity ^37,40^. In summary, the clinical syndromes of ATP1A3 mutations can be viewed as disorders of neuronal endurance and signal fidelity, where neurons that normally rely on *α3* can no longer sustain firing at the rates required for normal circuit function.

### Limitations and Future Directions

While our computational approach yielded clear predictions, it has inherent simplifications that must be acknowledged. First, the neuron model we used was based on a slowly adapting stretch receptor, yet with *α3* kinetics it exhibited high-frequency following characteristic of a rapidly adapting Ia afferent. Real muscle spindle primary endings have additional specializations – such as multiple spike initiation sites and modulatory fusimotor inputs – which we did not explicitly model. Incorporating these features could further extend the firing capabilities beyond the limit we observed. Conversely, our model neuron lacked certain channel dynamics (e.g. slow inactivation or activity-dependent ion concentration changes in restricted domains) that in real neurons might impose other limits on sustained firing. Thus, one future direction is to refine the model with more biophysical detail to see if the qualitative advantage of *α3* persists and to determine the true ceiling of 1:1 entrainment for *α3*-expressing neurons.

Another limitation is that we assumed an idealized substitution of pumps without other homeostatic adjustments. In vivo, neurons switching isoforms might also alter expression of ion channels or energy metabolism genes. Our simulations isolated pump kinetics as a variable; however, real neurons could partially compensate for a maladaptive pump by adjusting other properties (for example, an *α3*-deficient neuron might down-regulate certain K^+^ currents to become more excitable). Indeed, there are hints that chronic pump impairment can trigger complex changes in neurons ^35^. These feedbacks were beyond our scope but are important in understanding disease phenotypes. Experimentally, it remains challenging to swap or edit pump isoforms in adult neurons, but emerging genetic tools and pharmacological isoform-specific inhibitors ^41^ offer opportunities to test our predictions. For instance, selectively inhibiting *α3* in an intact preparation while recording high-frequency firing (as done with region-specific ouabain in mice ^41^) could validate the pump’s role in spike maintenance. On the flip side, overexpressing *α3* in normally *α1*-dominant neurons might be tested to see if it confers greater firing endurance.

Our discussion has largely focused on muscle spindle afferents and similar high-frequency neurons, but an open question is what role *α3* plays in other contexts such as nociceptors or cortical pyramidal cells. Most nociceptive dorsal root ganglion neurons predominantly express α1 pump^42^, consistent with their typically low firing rates or bursting patterns that do not demand long high-frequency trains. However, there is a subset of large-diameter DRG neurons (likely Aα/β fibers) that do express *α3*^10^. Intriguingly, some Aβ low-threshold mechanoreceptors can act as nociceptors after nerve injury (a phenomenon of A-fiber mediated pain) ^43^. One wonders if *α3* expression in certain mechanoreceptors might endow them with the capacity for high-frequency firing that could contribute to dysesthetic pain when disinhibited ^43^. Exploring isoform expression across sensory neuron subtypes as in ^3^ and relating it to their firing patterns is a worthwhile pursuit.

Finally, we emphasize the concept of neuron-specific pump tuning as a facet of excitability that has been under-appreciated. Just as neurons adjust ion channel profiles to shape action potentials and synaptic integration, they also “choose” a Na^+^ pump isoform to match their functional needs ^36,44^. This invites several speculative but intriguing questions. For example, could variation in pump isoform expression contribute to individual differences in sensory acuity or motor stamina? Could training or experience (e.g. endurance exercise, which elevates muscle spindle activity) induce shifts in *α3/α1* expression ratio in relevant neurons, thereby enhancing performance? There is some evidence that activity and growth factors can regulate Na^+^/K^+^-ATPase isoform levels in muscle and neurons ^5^. If so, pump isoforms might be plastic components in the nervous system’s long-term adaptation to functional demands. On the other hand, because *α3* is so critical, neurons may have limited ability to compensate if it is lost – explaining why de novo ATP1A3 mutations have devastating effects rather than benign phenotypic variation. Going forward, interdisciplinary studies combining detailed biophysical modeling, genetic manipulation of pump isoforms, and behavioral physiology (such as measuring vibration perception or motor endurance in animal models) will help to fully elucidate the role of Na^+^/K^+^-ATPase isoform kinetics in neural function. Our current findings provide a mechanistic foundation, showing that the *α3* isoform is not merely an evolutionary redundancy, but a key enabler of neuronal high-performance – allowing certain neurons to fire fast, long, and faithfully to meet the demands of their specialized roles.

## Methods

### Gestrelius model

The receptor model was adapted from ^45^ as follow

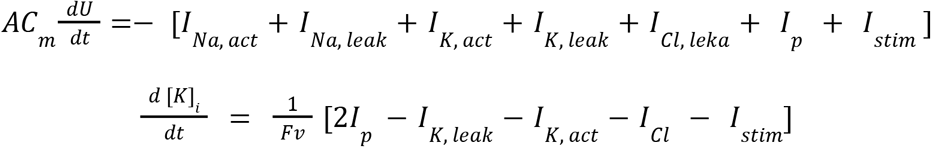

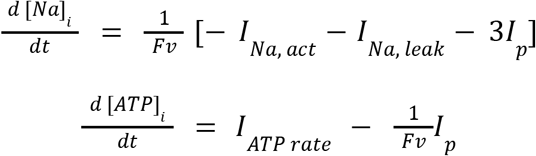

where U is membrane potential; *I*_*Na, act*_, *I*_*K, act*_ are the active sodium and potassium currents; *I*_*K, leak*,_*I*_*Na*,*leak*_, *I*_*Na*,*leak*_ are the leak currents for potassium, sodium and calcium; *I*_*p*_ is the current generated by the Na-K-ATPaset, *I*_*stim*_ is the external stimulation current; [*K*] _*i*_, [*Na*]_*i*_ and [*APTP*] _*i*_ are the intracellular concentrations of potassium, sodium, and ATP respectively; A is the projected membrane, *C*_*m*_ is the membrane capacitance, F is the Faraday constant. Equations for the currents are:

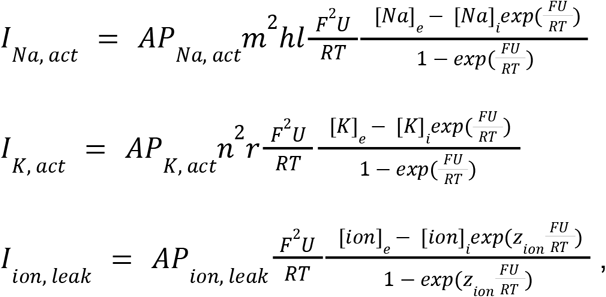

where *I* _*ion, leak*_ represents the leak current for potassium, sodium and calcium, with ‘ion’ denoting for *Na*^+^, *K*^+^ and *Cl*^−^, respectively. R is the universal gas constant; T is the absolute temperature, intra- and extra-cellular concentration of ions are denoted as [] _*i*_ and []_*e*_, respectively. P is the maximum permeability. m, h, l, n and r stand for channel gating variables. The equations and parameter values for these variables are taken from ^14^:

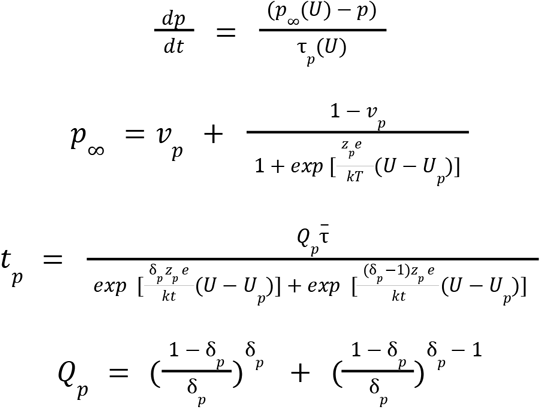

Here, *p* denotes one of the five gating variable, *v*_*p*_ is the minimum value of the corresponding channel permeability, *z*_*p*_ is the effective valency of the gating particle, *e* is the electron charge, *k* is the Boltzmann constant, δ_*p*_ is the degree of energy barrier asymmetry varying within, *U*_*p*_ is the membrane potential at which half of the corresponding gating channels are open, 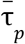 is the maximum value of τ_*p*_. The parameters are used in this work taken for the (SAO) slowly adapting organ model.

The Na-K-ATPase current for *a* -isoform presented as follow:

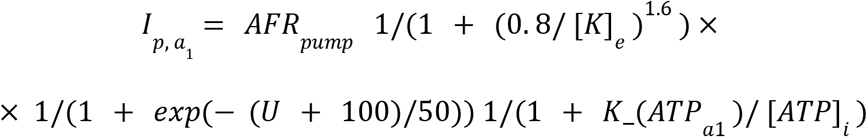

and for the *a*_3_ -isoform, it’s given by:

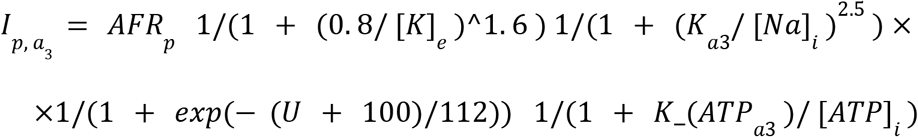

where *R*_*p*_ is the maximum pump current. The Hill coefficients for potassium and sodium were taken from ^8^, the half activation constant for the *a*_1_ -isoform 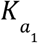 is 10 mM, and for the *a*_3_ -isoform 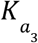 is 30 mM ^1^. The half-activation constant for ATP 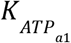 is 0.4 mM for the α1 isoform 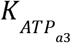 and 0.2 mM for the α3 isoform as described in the Michaelis–Menten equation ^7^. The slope and half activation voltage for voltage dependency were selected based on the from ^8^.

The surface area of the modeled DRG neuron was set to 10000 μ*m*^2^ assuming a spherical geometry, which corresponds to cell volume of 9.4 μ*m*^3^. Given this data, the value of the Na-K-ATPase current at rest for pump with *a*_1_ -isoform is 800 pA ^17^. Cell membrane capacitance is 1 μ*F*/*cm*^2^.

Membrane permeability constants were derived from the membrane conductance of DRG mouse neurons of the same size as noted above ^15^, calculated as the partial derivative at the reverse potential for sodium and according to the following equation:

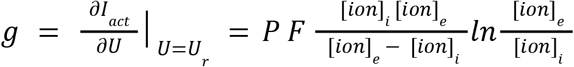

Constants for the leak permeabilities of sodium, potassium, and calcium, as well as the maximum pump current, were calculated so that the model at rest maintains: a membrane potential of –0.065 mV, an intracellular sodium concentration of 7 mM, an intracellular potassium concentration of 147 mM, and a Na-K-ATPase current of 800 pA while simultaneously preserving the potassium-to-calcium permeability ratio of 10:3. Extracellular concentrations were set to 145 mM for sodium, 3.5 mM for potassium, and 15 mM for calcium, while the intracellular calcium concentration was 15 mM. ATP levels were assumed constant during this procedure. Subsequently, the ATP synthesis rate was adjusted to maintain intracellular ATP at a physiological level 5 mM ^16^. The parameters obtained from this procedure were then used for other isoforms of Na-K-ATPase and their modifications. All simulations were initiated once the model reached its resting state.

### Dorsal root ganglion isolation and primary culture

Adult male Harlan Sprague–Dawley rats (250–350 g; Harlan Sprague–Dawley, Indianapolis, IN) were euthanized by cervical dislocation under ether anesthesia. Eight to twelve lumbar dorsal root ganglia (DRG) were dissected and transferred into a bicarbonate-buffered saline containing (in mM): 130 NaCl, 2 CaCl_2_, 2 MgSO_4_, 3 KCl, 1.25 KH_2_PO_4_, 10 D-glucose, and 26 NaHCO_3_; pH 7.3, 310 mOsm. Ganglia were cleared of connective tissue and rinsed twice in the same buffer lacking CaCl_2_. They were then incubated in the same buffer supplemented with 0.5% collagenase (Worthington Labs, Lakewood, NJ) for 30 min at 37 °C (water bath). After digestion, tissues were rinsed twice with the Ca^2+^-free buffer and twice with a HEPES-buffered saline containing (in mM): 156 NaCl, 2 CaCl_2_, 2 MgSO_4_, 3 KCl, 1.25 KH_2_PO_4_, 10 D-glucose, and 7.5 HEPES. Following a 10-min incubation at 37 °C, tissue was triturated to dissociate cells. Dissociated cells were plated on 60-mm tissue-culture dishes (Falcon, Becton-Dickenson, Lincoln Park, NJ) and maintained in Medium 199 (HEPES modification; Gibco-BRL Laboratories, Gaithersburg, MD) supplemented with 1% penicillin–streptomycin and 10% fetal bovine serum. Cultures were kept in a water-jacketed incubator at 37 °C with 5% CO_2_ until use. Recordings of DRG neuron spike responses were performed in standard physiological saline without specific blockers, as in the pump-current measurements.

### Patch-clamp recordings

Experiments were performed on freshly isolated neurons or on neurons maintained in short-term culture (1–2 days). Whole-cell recordings were obtained with standard techniques using a DAGAN 3900 patch-clamp amplifier with whole-cell extender (3911A; Dagan Corp., Minneapolis, MN). Neurons were identified by soma size: for cells with diameters <25 µm, 10-ms steps from a holding potential of −60 mV to 0 mV were applied to verify a voltage-dependent Na^+^ current; larger cells were assumed to be neurons. Voltage commands and data acquisition were controlled by pCLAMP software with an AxoLab 1100 interface (Axon Instruments, Burlingame, CA). Patch electrodes were pulled from Corning 7052 glass (Garner Glass Co., Claremont, CA) and fire-polished to a 1–5 µm tip diameter (1–5 MΩ when filled). Liquid-junction potentials were nulled immediately before seal formation. Holding current (I_h) at a holding potential (V_h) of −40 mV was continuously monitored on a chart recorder (The Recorder Co., San Marcos, TX).

## Data Availability

Code and model files are available at: https://github.com/Reshetnikoff/Na-K_ATPase_model.git

## Author contribution

M.G. Dobretsov is supported by IEPB RAS government basic research program #075-00264-24-00.

## Notes

https://github.com/Reshetnikoff/Na-K_ATPase_model#

### Competing Interest Statement

The authors have declared no competing interest.

